# Benchmarking PSM identification tools for single cell proteomics

**DOI:** 10.1101/2021.08.17.456676

**Authors:** Daisha Van Der Watt, Hannah Boekweg, Thy Truong, Amanda J Guise, Edward D Plowey, Ryan T Kelly, Samuel H Payne

## Abstract

Single cell proteomics is an emerging sub-field within proteomics with the potential to revolutionize our understanding of cellular heterogeneity and interactions. Recent efforts have largely focused on technological advancements in sample preparation, chromatography and instrumentation to enable measuring proteins present in these ultra-limited samples. Although advancements in data acquisition have rapidly improved our ability to analyze single cells, the software pipelines used in data analysis were originally written for traditional bulk samples and their performance on single cell data has not been investigated. We benchmarked five popular peptide identification tools on single cell proteomics data. We found that MetaMorpheus achieved the greatest number of peptide spectrum matches at a 1% false discovery rate. Depending on the tool, we also find that post processing machine learning can improve spectrum identification results by up to ∼40%. Although rescoring leads to a greater number of peptide spectrum matches, these new results typically are generated by 3rd party tools and have no way of being utilized by the primary pipeline for quantification. Exploration of novel metrics for machine learning algorithms will continue to improve performance.

## Introduction

Single cell proteomics has gained interest in recent years because it directly interrogates cellular heterogeneity. Biological tissues are made up of a complex mix of specialized cells, and the analysis of a homogenized sample prevents the characterization and understanding of the individual cell types and their interactions^1,2^. Analysis of bulk samples has previously been the standard because mass spectrometry (MS) has historically required thousands to millions of cells to achieve good quality data. However, recent improvements in experimental methods and instrumentation have enabled single cell proteomics^3–5^.

Although data generation has greatly improved to support single cell proteomics, algorithms for data analysis are still largely derived from bulk proteomics data. Algorithm development is a popular area in computational mass spectrometry, and specialized tools exist for bottom-up, top-down, glycomics, metabolomics and many other applications. While several tools have been developed for post-identification statistical data wrangling of single cell proteomics data^6,7^, the critical and first step of data analysis — spectrum identification — has not yet been evaluated. Here, we benchmark five popular tools for peptide identification on single cell-sized aliquots of a commercial HeLa digest. We also demonstrate that a significant boost in performance is provided by machine learning re-scoring for trace samples.

## Results

As peptide identification software tools were designed and optimized for spectra from traditional bulk sample proteomics, it is important to understand how they perform for spectra from ultra-small samples like single cells. The spectra used in this evaluation were from 12 instrument runs: six from a 2 ng sample which approximates 10 cells and six from a 0.2 ng sample which approximates 1 cell (see Methods). We benchmark 5 commonly used peptide identification tools: MetaMorpheus^8^, MsFragger^9^, MSGF+^10^, MaxQuant^11^, and Proteome Discoverer^12^.

### Tool Benchmarks

Peptide identification algorithms assign each peptide/spectrum match (PSM) a confidence score. Unfortunately, there is no universally employed statistic in proteomics to express the probability of a PSM being correct^13^. The five tools in our study use three different statistical metrics: q-values, posterior error probabilities (PEP) and the PeptideProphet Score. For our benchmarking, we chose to evaluate the number of peptides detected by each tool using its own native statistical score. Search parameters and settings were carefully chosen to be as similar as possible (see Methods), and when possible were adjusted according to recommendations from the developers of each tool.

The spectral files were processed by each tool and the number of PSMs passing a 1% cutoff were totaled. Although there is some variation in the number of PSMs between files, each tool’s performance is consistent across the entire dataset, with MetaMorpheus identifying the most PSMs per file (Figure 1A). For 2 ng data, MetaMorpheus typically identified 9% more PSMs than MSGF+, 16% more than MSFragger, and 34% more than MaxQuant. We were concerned that the significantly fewer PSMs for MaxQuant might be attributed to the use of PEP, which is fundamentally different from q-value. Fortunately, MetaMorpheus natively computes both PEP and q-value, thus we were able to directly compare the MaxQuant results to MetaMorpheus. The trends identified in the 2ng samples were also observed in the 0.2 ng samples (Figure 1B).

**Figure 1.**
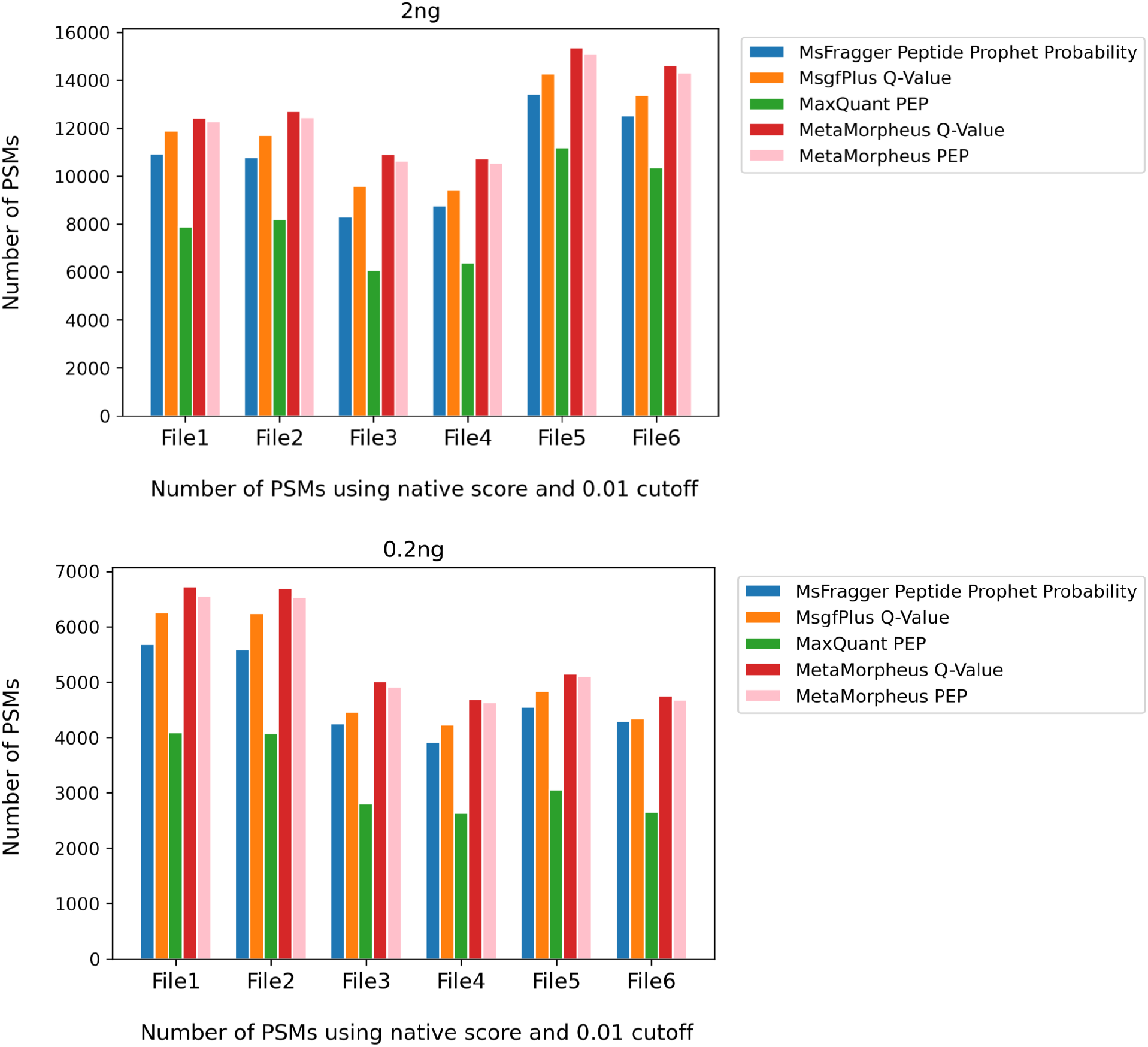
Number of PSMs identified for each tool. The number of PSMs identified with a high confidence score (FDR <= 0.01) for each peptide identification tool, using the tool’s native probability score. MSGF+ uses a Q-Value score, whereas MaxQuant uses a PEP score, and MsFragger uses a Peptide Prophet Probability. MetaMorpheus provides both PEP and Q-Value and both are included to provide a comparison for PEP and Q-Value scores. MetaMorpheus consistently has the most PSMs across all files for both 2 ng and 0.2 ng. MSGF+ has the second most PSMs and MSFragger has the third. MaxQuant has the fewest PSMs counted.

### Machine Learning Re-scoring

Machine learning can be used to improve performance by optimizing the weight for features within a scoring function^14^. Semi-supervised machine learning tools like Percolator^15^ and mokapot^16^ re-evaluate feature weights to discriminate between correct and decoy PSMs. These tools utilize a variety of input features, which typically arise from a peptide identification algorithm and include metrics such as number of *y* ion peaks, % spectrum intensity within the *b*/*y* ladder, precursor ppm error, etc. A significant benefit of using a re-scoring tool is that identifications from different algorithms are all ranked and scored with a consistent statistic.

We use mokapot for post-hoc re-scoring of each tool’s PSM output. We also included PSM identifications from ProteomeDiscoverer, which automatically uses Percolator to re-score its results. After re-scoring, MetaMorpheus was still the most effective algorithm (Figure 2), typically scoring 10% more than other tools. As each algorithm has a unique set of features describing the PSMs, mokapot’s performance varies substantially between algorithms. For example, with 2 ng data, MetaMorpheus saw an 12.0% increase, MsFragger increased by 18.3%, and MaxQuant increased by 40.7%. MSGF+ had a limited set of features in the PSM table, and therefore, the changes in post-hoc re-scoring were negligible. Similar improvements for each tool were seen for the 0.2 ng samples.

**Figure 2.**
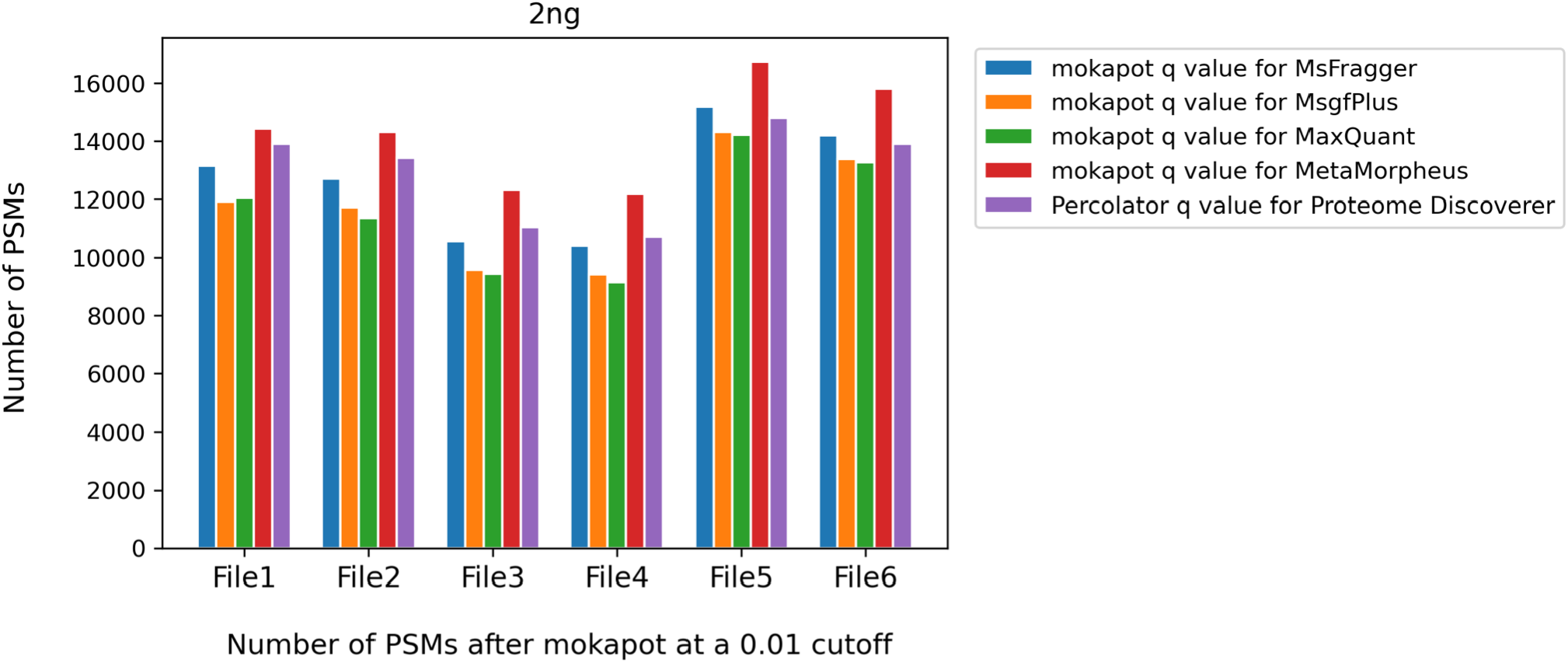
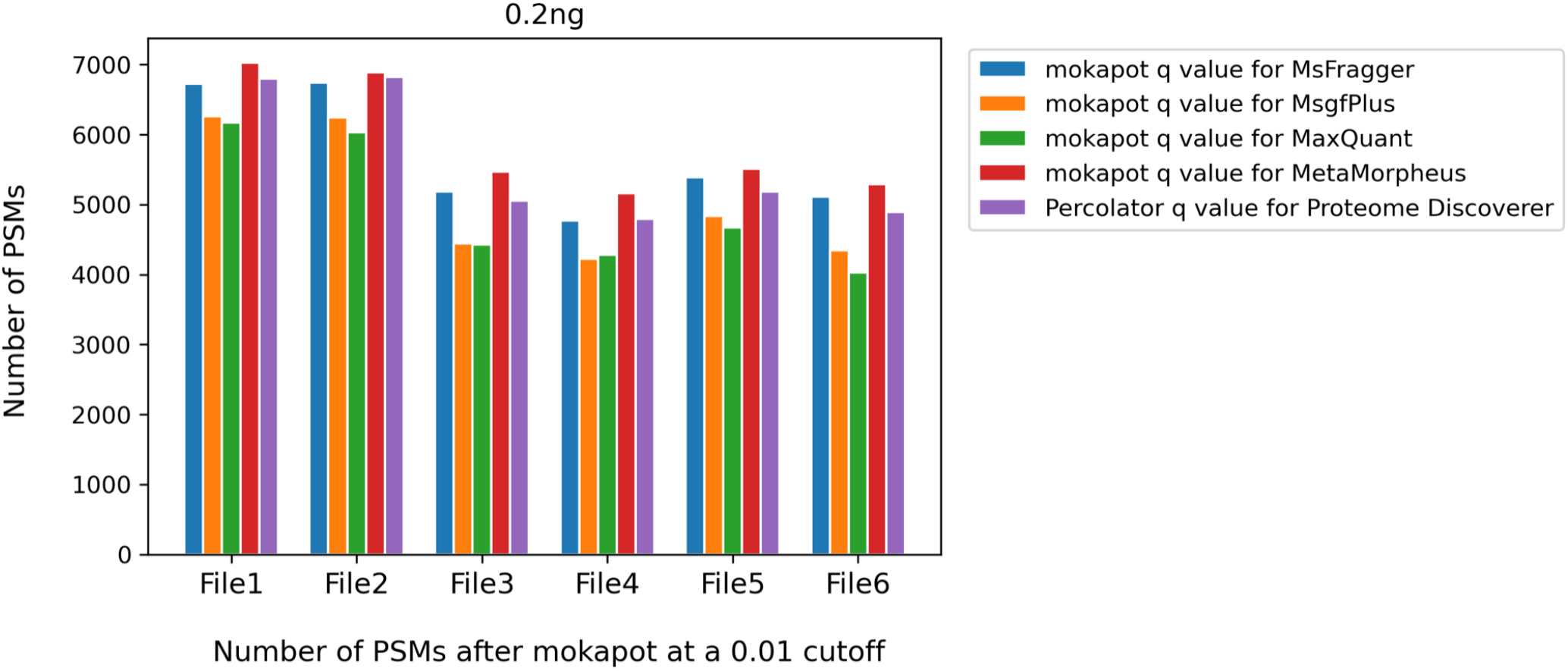
improved PSM identification with machine learning re-scoring. Figure 2 has an additional algorithm, Proteome Discoverer. This algorithm synchronously runs Percolator, a rescoring program similar to mokapot, thus, we do not have the native data from Proteome Discoverer. The rescored results given are more similar to the rescored data given in Figure 2 than to the native data in Figure 1. Similar to Figure 1, MetaMorpheus continually scores the highest number of PSMs under the cutoff. For 2 ng data, MetaMorpheus typically identified 18.0% more PSMs than MSGF+, 11.0% more than MSFragger, 19.3% more than MaxQuant, and 14.0% more than Proteome Discoverer. The trends in 0.2 ng data are very similar; MetaMorpheus typically identified 14.7% more PSMs than MSGF+, 4.3% more than MSFragger, 16.6% more than MaxQuant, and 11.1% more than Proteome Discoverer.

### New Features for Machine Learning Re-scoring

In the previous section, the only features given to mokapot were those that were present in the PSM output from each tool. However, given the varied improvement observed above, we were curious if adding new features to a program’s output could continue to improve PSM identifications. As MetaMorpheus outperformed other algorithms and it had an extensive set of features in the PSM identification file, we specifically investigated whether the addition of new features could improve its performance. We identified and tested three new features: the longest series of consecutive *y* peaks, the percent of *y* peaks in a consecutive series, and the absolute difference between predicted retention time and actual retention time (see Methods).

After adding these new features to the existing MetaMorpheus data, we re-evaluated the improvement due to mokapot re-scoring. When comparing mokapot with the original output of MetaMorpheus to mokapot with extra features, peptide identifications increased. Without rescoring, MetaMorpheus identified 12,446 PSMs at qvalue < 0.01. With mokapot rescoring, an average of 14,420 PSM are identified. When additional features are added, mokapot rescoring produced an average of 14,785 PSMs - an extra 2.9% increase in PSMs (t-test, p< 0.05).

### Living Benchmark

As single cell proteomics is a rapidly developing field, we expect continued development in new and improved algorithms. To facilitate future comparisons with these tools, we wrote a living benchmarking tool that allows users to compare any tool to the results of this publication. The mass spectral files for the 12 datasets used here are available on MassIVE (see Methods). To compare a new algorithm’s PSMs to the results in Figure 1, follow the instructions within our GitHub repository (https://github.com/PayneLab/SingleCellBenchMark/blob/main/Living_BenchMarking/).

## Discussion

We benchmarked multiple popular peptide identification tools for their performance on single cell proteomics data. We note that performance is highly variable between tools and that the result of each tool improved with machine learning post processing of PSM results. In addition, we demonstrate that adding new features further improved performance. Therefore, we suggest that algorithm developers continue to explore metrics which differentiate target and decoy spectra.

Although PSM and peptide identification is improved with machine learning re-scoring, it is often not part of routine use within the proteomics community. Many software tools integrate both identification and quantification into a single workflow and lack a way for a 3rd party to integrate their improved peptide identification results back into the workflow. Given the 10-30% increase in PSM identifications, software tools and pipelines should become more flexible and allow for 3rd party interaction. Increasing the list of confidently identified PSMs would improve overall protein identification and quantification.

## Methods

All software used in the data generation and analysis is publicly available in our GitHub repository, https://github.com/PayneLab/SingleCellBenchMark.

### Datasets

Twelve mass spectrometry runs of the Pierce HeLa digest were used in this manuscript, consisting of six runs with 2 ng of peptide input and six runs of 0.2 ng of peptide input. These datasets have been previously described and are freely available at MassIVE (MSV000087689).

### Algorithm Configuration and Execution

For each of the algorithms, we attempted to make the runtime parameters as similar as possible. We also contacted algorithm developers and tool maintainers for specific suggestions for single cell data. General parameters include: peptide length 8-30; fixed modification of carbamidomethyl, and a variable modification of oxidation; two variable modifications per peptide; precursor and fragment mass tolerance 20 ppm. Specific parameter files for each tool are uploaded to our GitHub repository (∼/Parameters).

#### PSM output

The primary comparison in our manuscript is the number of PSMs at a given statistical confidence. The 12 files were run through each algorithm and the output is stored in GitHub at “∼/data/”. For each tool we specify the output file used. <u>MetaMorpheus:</u> For figure 1, we used the psmtsv files and QValue and PEP. However, following the developers recommendation, the pin file was used for machine learning re-calibration. <u>MaxQuant:</u> We used the msms.txt file and PEP. <u>MsFragger:</u> PSM identifications were taken from the psm.tsv.gz file. As PeptideProphet Probability is calculated for the likelihood a PSM is correct (instead of the more common probability that it is false), we internally converted these to 1-PeptideProphetProbability. <u>MSGF+:</u> mzID files were converted to a tab format using the developer’s recommentations at https://msgfplus.github.io/msgfplus/MzidToTsv.html. The QValue column was used for statistical confidence. <u>Proteome Discoverer:</u> Proteome Discoverer automatically runs the Percolator algorithm on all PSM identifications; a cutoff of Percolator q-Value < 0.01 was used for data included in Figure 2.

**Figure 1.** Decoys and duplicate scans were filtered out, and the PSMs were counted and aggregated across all files. The exact code for making Figure 1 can be found in the file “∼/makeFigure1.ipynb”.

### Rescoring with Mokapot

Before running PSM results through mokapot, each file was cleaned by trimming multiple PSMs per scan number, keeping the one with the best score. Feature columns with missing values were removed, as mokapot requires a complete data matrix. MetaMorpheus produces a file that is specifically formatted post-hoc rescoring. However, for all the other algorithms the feature columns had to be hand picked as the following:

<u>MSGF+ :</u> Charge, IsotopeError, PrecursorError(ppm), DeNovoScore, MSGFScore, and SpecEValue.

<u>MsFragger:</u> Charge, Peptide Length, Retention, Delta Mass, Expectation, Hyperscore, Nextscore, Number of Enzymatic Termini, Number of Missed Cleavages, and Intensity.

<u>MaxQuant:</u> Charge, Precursor Intensity, Score, Length, Missed cleavages, m/z, Mass, Retention time, and Delta score.

The exact code for running the files through mokapot is found in our GitHub in the “∼/MokaPot/” directory. Each tool has a notebook that is named “MokaPot_[tool name].ipynb. In our GitHub repository, the re-scored PSMs can be found at “∼MokaPot/MokaPot_Output/”. For Figure 2, the number of PSMs with mokapot q-value < 0.01 (or the Percolator q-Value for ProteomeDiscoverer) were counted and totalled. The exact code for making Figure 2 can be found in the GitHub repository at “∼/make_Figure2.ipynb”. The calculations for the average increase for each tool’s results after mokapot rescoring can be found inside “∼/Avg_PSM_count_change”.

### Adding New Features

Three new columns were added onto the PSM identifications of MetaMorpheus as additional features for mokapot to use in its rescoring: number of consecutive *y* peaks, the percent of annotated peaks in a consecutive series, and the absolute difference between predicted retention time and actual retention time^17^. These new features were joined with the MetaMorpheus features (pin file), and run through mokapot. The cross-validation partitioning of PSMs within mokapot is randomly determined, and therefore the results are very similar but not identical each time the program is run. To account for this variability, we report the average of 25 independent mokapot rescorings. The code to add the new features and calculations can be found in the repository at “∼/Optimization/” and “∼/updated_features.ipynb”.

